# Resolving drug effects in patient-derived cancer cells links organoid responses to genome alterations

**DOI:** 10.1101/124446

**Authors:** Julia Neugebauer, Franziska M. Zickgraf, Jeongbin Park, Steve Wagner, Xiaoqi Jiang, Katharina Jechow, Kortine Kleinheinz, Umut H. Toprak, Marc A. Schneider, Michael Meister, Saskia Spaich, Marc Sütterlin, Matthias Schlesner, Andreas Trumpp, Martin Sprick, Roland Eils, Christian Conrad

## Abstract

Cancer drug screening in patient-derived cells holds great promise for personalized oncology and drug discovery but lacks standardization. Whether cells are cultured as conventional monolayer or advanced organoid cultures influences drug effects and thereby drug selection and clinical success. To precisely compare drug profiles in differently cultured primary cells, we developed *DeathPro*, an automated microscopy-based assay to resolve drug-induced cell death and proliferation inhibition. Using *DeathPro*, we screened cells from ovarian cancer patients in monolayer or organoid culture with clinically relevant drugs. Drug-induced growth arrest and efficacy of cytostatic drugs differed between the two culture systems. Interestingly, drug effects in organoids were more diverse and had lower therapeutic potential. Genomic analysis revealed novel links between drug sensitivity and DNA repair deficiency in organoids that were undetectable in monolayers. Thus, our results highlight the dependency of cytostatic drugs and pharmacogenomic associations on culture systems, and guide culture selection for drug tests.

## Introduction

Cell-based assays are a key tool in basic research and drug discovery, and are increasingly used in personalized oncology. In the last years, numerous anticancer therapeutics developed from standard cell line screens in conventional 2D culture failed in clinical studies^1^. As a result, standard treatment and overall survival of advanced cancers like ovarian cancer (OC) has not changed for decades^2^. To allow personalized therapy and improve drug development, new patient-derived models such as organoids^3–5^ and patient-derived xenografts^6–8^ that recapitulate the heterogeneity and intrinsic drug sensitivity of the original tumour have started to replace the popular cancer cell lines. Patient-derived organoids may be grown as 3D cultures on hydrogels like Matrigel that mimic the extracellular matrix. Compared to 2D cell cultures, they have emerged as near-physiological models reflecting the gene expression, differentiation and structure of the primary tissue^9^. Nevertheless, due to increased workload, higher costs and the current lack of 3D assay methods, most drug screens are still performed in less physiological 2D cultures^10^. Initial studies in ovarian and breast cancer showed that cells cultured as cell aggregates are less sensitive to drugs than in monolayer culture^11,12^. The culture format thus shapes cellular drug responses and defines the translational power of a drug assay. However, this dependency cannot be studied in detail with widely-used, unspecific viability assays that measure metabolic activity or cellular ATP as surrogate markers. Such assays show limited reproducibility and do not resolve actual drug effects of high therapeutic interest such as cell death and growth arrest^3,13^. Instead, recent advances in automated microscopy enable more sophisticated assays that can de-convolve drug effects in different culture formats.

Here, we systematically compare drug effects in organoid and standard 2D culture using *DeathPro*, a confocal microscopy based assay and image processing workflow to simultaneously study cell death and growth arrest in patient-derived material over time. Using *DeathPro*, we screened cells from nine high-grade serous OC patients with clinically relevant drugs and found that growth arrest and the efficacy of cytostatic drugs notably depends on the culture type. Remarkably, patient-specific genomic alterations correlated with drug effects observed in organoids, but not in 2D cell monolayers. Hence, combining refined assays like *DeathPro* with advanced models like cancer organoids could enhance drug screening in the context of personalized oncology and pharmacogenomics.

## Results

### De-convolving drug-induced cell death and proliferation inhibition

To resolve drug effects in patient cells and organoids, we developed an automated live cell assay and quantification workflow, which de-convolves drug-induced **death** and **pro**liferation inhibition over time (*DeathPro*) (Fig. 1). To this end, cells were stained with Hoechst and counterstained with propidium iodide (PI) for dead cells and analysed at consecutive time points by confocal microscopy. To accurately quantify cell growth for each condition^14^, cells were imaged at the start and end of the drug treatment at the same position. For high-throughput image analysis, we built an adaptable visual programming workflow that encompasses adaptive sequential thresholding and outlier filtering strategies to cope with heterogeneous cell morphologies and dye intensities. In the workflow, total areas covered by dead cells (PI-stained) and all cells (Hoechst or PI-stained) were determined from confocal images and used to calculate LD50 values and area under curve values for cell death (AUCd) and proliferation inhibition (AUCpi) (Fig. 1b).

**Figure 1:**
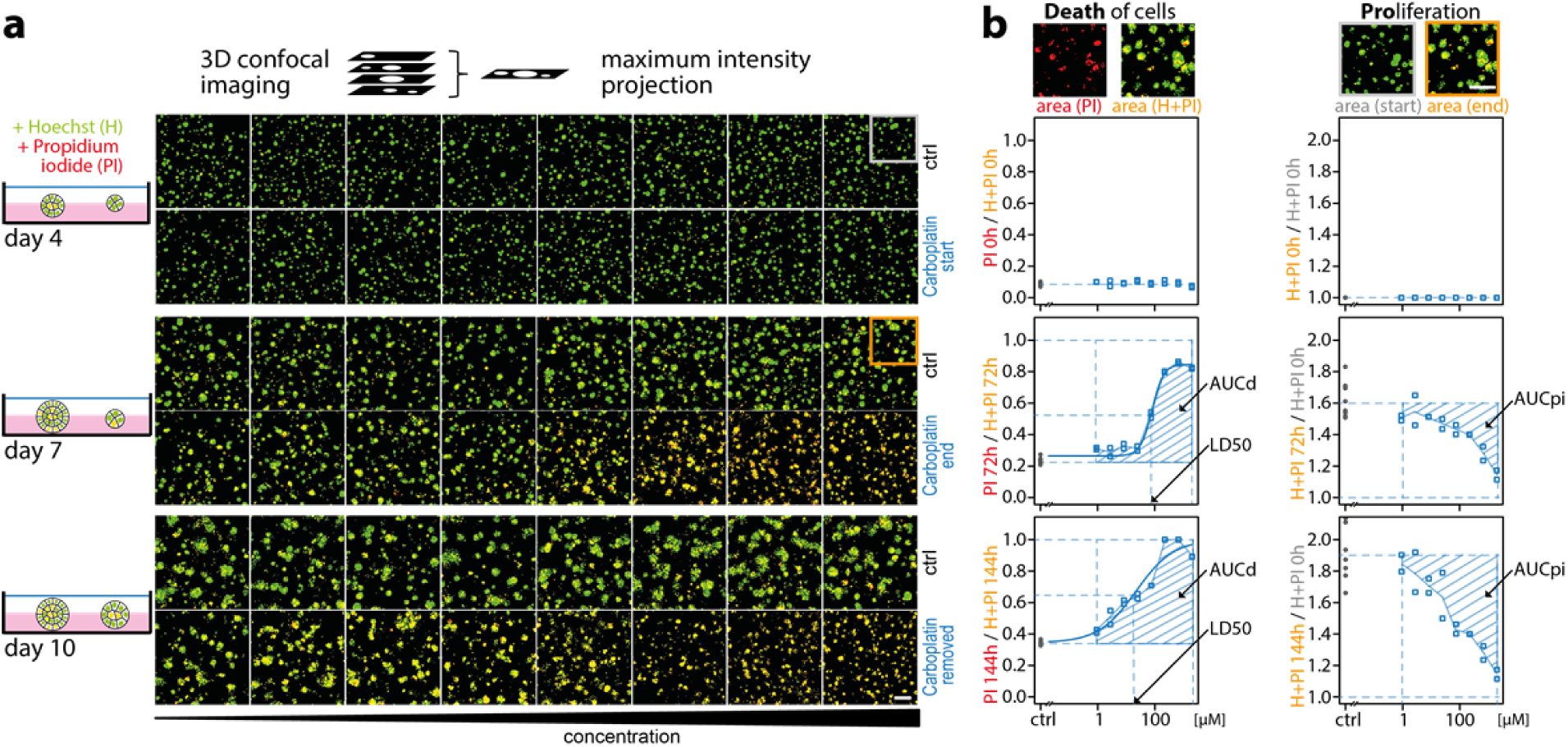
Drug-induced cell death and proliferation inhibition can be quantified from serial confocal images. (**a**) Schematic overview of drug testing in organoid culture with the DeathPro assay. Cells are grown on Matrigel for 4 days, stained with Hoechst (H) and Propidium iodide (PI) and imaged at day 4, day 7 and day 10. Image gallery exemplifies OC12 organoid growth and cell death at start (day 4) and end of carboplatin treatment (day 7) and after carboplatin removal (day 10) using eight carboplatin concentrations or drug-free medium (ctrl). Confocal images are reduced to maximum intensity projections and binary images of merged Hoechst (green) and PI (red) channel are shown. (**b**) Image analysis for the DeathPro assay is based on area measurements in Hoechst and PI channels, and calculation of LD50, AUCd and AUCpi values to describe cell death and growth arrest. Drug response curve fitting and AUC values are illustrated for OC12 at 0 h and 72 h time points depicted in (a). Grey and orange boxes in (a) correspond to the magnifications in (b). Scale bar is 200 μm.

The *DeathPro* assay and workflow reliably resolved carboplatin-induced cell death and proliferation inhibition in OC patient-derived organoids (Fig. 1). In addition, we resolved drug effects in lung cancer organoids (Supplementary Fig. 1) to verify that the *DeathPro* workflow can be applied to patient cells from different cancer entities.

By using live cell dyes, patient cells or organoids can be directly used for screening and do not have to be genetically modified to express fluorescent proteins. To exclude the possibility that either dye alters cell behaviour, we tested their effect on ovarian organoids. Hoechst and PI did not affect organoid growth but increased cell death (Supplementary Fig. 2), which is accounted for in AUCd measurements by normalization to the untreated control (Fig. 1b). Imaging OC12 organoids only at the end or additionally at the beginning of the drug treatment did not alter organoid growth or cell death (Supplementary Fig. 2).

### Drug-induced growth arrest in ovarian cancer patient cells is culture-dependent

To systematically assess culture-type influence on patient cell responses, we used the *DeathPro* assay to screen patient-derived OC cell lines (PDCLs) cultured in standard 2D culture or as cancer organoids. PDCLs were established from metastatic serous ovarian cancers (FIGO stage IIIc-IV, Supplementary Table 1, Fig. 2a). Additionally, we included Human Ovarian Surface Epithelial cells (HOSEpiC) to assess potential side-effects such as cytotoxicity in normal cells. Seeded on Matrigel, HOSEpiC and PDCLs developed into spheres or morphologically diverse ‘cancer organoids’ (Fig. 2b), respectively, with the latter retaining expression of the tumour markers CA-125 and WT1 (Supplementary Fig. 3).

**Figure 2:**
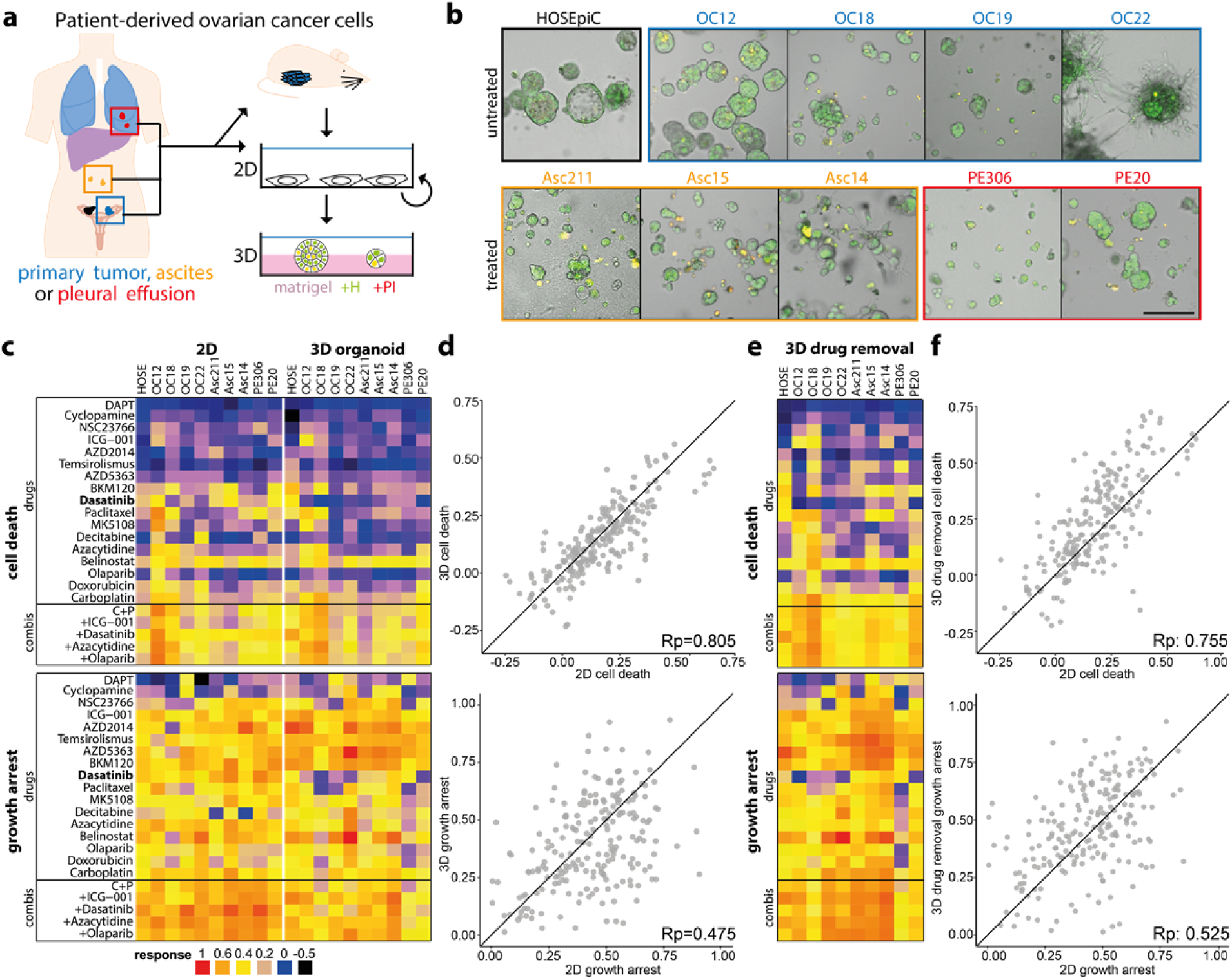
Culture type shapes drug-induced growth arrest in ovarian cancer patient cells. (**a**) Simplified overview of generation and cultivation of patient-derived ovarian cancer cell lines (PDCLs) from different sites (OC: primary tumour, Asc: ascites, PE: pleural effusion). Patient material was taken directly into 2D culture or amplified by xenografting into mice. PDCLs are cultured in 2D culture but can be grown as ovarian cancer organoids on Matrigel (**b**) Morphology of ovarian cancer organoids and normal ovarian epithelial cells (HOSEpiC) on Matrigel seven days after seeding. Green (Hoechst) and red (PI) channels are merged. (**c**) Drug responses (cell death: AUCd, growth arrest: AUCpi) measured with *DeathPro* assay after 72h drug treatment in patient cells cultured as monolayers (2D) or ovarian cancer organoids (3D). (**d**) Comparison of drug-induced cell death (AUCd) and growth arrest (AUCpi) in 2D vs 3D. (**e**) Drug responses measured in ovarian cancer organoids (3D) after 72h drug treatment followed by 72h drug removal. (**f**) Comparison of drug-induced cell death and growth arrest in 2D vs 3D after drug removal. All values shown are means of two independent biological replicates. HOSE=HOSEpiC, Rp = Pearsons correlation coefficient. C+P = Carboplatin + Paclitaxel

OC organoids or 2D cultured PDCLs were screened twice for 22 drugs or drug combinations (Supplementary Table 2) currently used or under investigation for treatment of OC. LD50 and cell death (AUCd) values were highly reproducible across all drugs and patients in 2D and organoid culture (Pearson correlation 0.86-0.97, Supplementary Fig. 4a), whereas growth arrest (AUCpi) showed slightly lower correlation (Pearson correlation 0.67-0.76, Supplementary Fig. 4b).

Based on the *DeathPro* results, we compared all drug effects determined in OC patient cells between 2D and 3D culture (Fig. 2c). In both screens, drugs induced more growth arrest than cell death (Fig. 2d). Due to low drug-induced cell death, LD50 values could not be determined in 20-30% of all conditions (Supplementary Fig. 5a). After 72h drug treatment, cell death was slightly lower in organoids than in 2D cultures (Fig. 2c, Supplementary Fig. 5b). Surprisingly, death upon drug treatment strongly correlated in 2D and 3D culture whereas drug-induced growth-arrest varied greatly with culture type (Fig 2d, Pearson correlation 0.85 vs 0.475). Since drug-induced cell death was growth-dependent and organoids grew slowly compared to cells in 2D culture (Supplementary Fig. 3c), we measured organoid responses a second time after drug removal in 3D (Fig 2e, Supplementary Fig. 6a). After wash out, drug effects were increased in most patient organoids (Supplementary Fig. 6a-c) but cytotoxicity levels still resembled those in 2D culture (Fig. 2f, Pearson correlation 0.755). Likewise, LD50s measured in 3D culture before and after drug removal highly correlated with LD50s in 2D culture (Supplementary Fig. 5c, d, Pearson correlation 0.872, 0.822). In contrast, growth inhibition again differed after drug removal (Fig. 2f, Pearson correlation 0.525). Overall, growth arrest was the major drug effect in OC cells and was culture-type dependent, whereas cell death was similar between culture types.

### Efficacy of cytostatic drugs depends on culture type

As the culture type affected growth arrest, the efficacy of cytostatic drugs that do not induce cell death should be culture-type dependent as well. Thus, we compared the efficacy of drugs in our panel by summarizing drug response parameters (LD50, AUCd and AUCpi) into a single efficacy measure, and clustering drugs on this basis. In 2D and 3D culture, three clusters arose based on differential cytotoxicity: (i) drugs effectively inducing cell death and growth inhibition (red cluster), (ii) medium cytotoxic drugs (yellow cluster) and (iii) ineffective drugs (blue cluster, Fig. 3a, b). Clustering revealed that the most effective treatments (red cluster, Fig. 3a, b) in both screens comprised belinostat, BKM120, the first-line therapeutic carboplatin and all combinations thereof. Paclitaxel, which forms part of the current first-line therapy for OC, was not among the most effective treatments tested due to its low toxicity in most patient cells (Fig. 2c, e). Moreover, its combination with carboplatin performed no better than carboplatin alone in 2D and 3D (Supplementary Fig. 6c, d).

**Figure 3:**
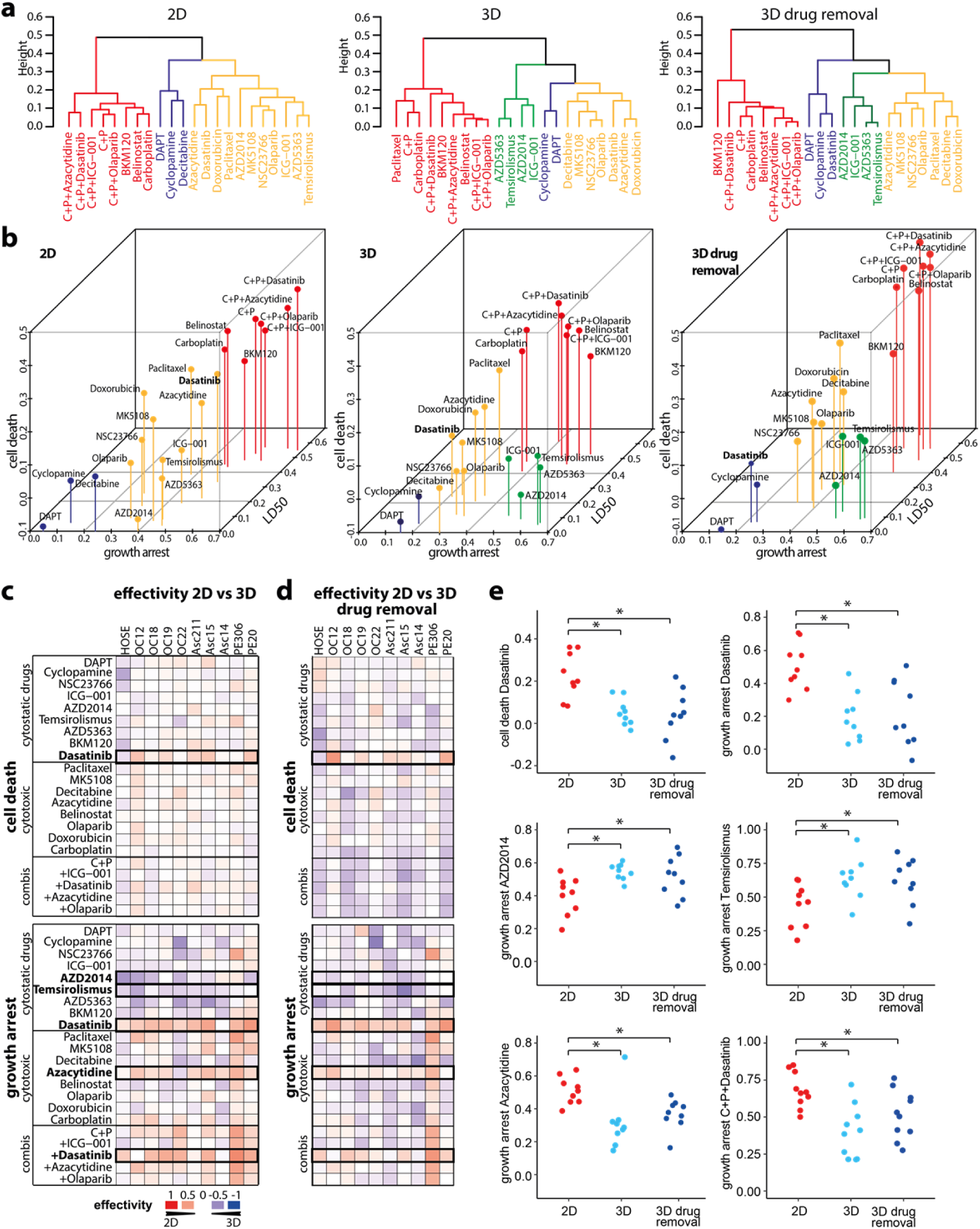
Culture type determines effectivity of targeted drugs like SRC inhibitor dasatinib and mTOR inhibitors AZD2014 and temsirolismus. (**a**) Hierarchical clustering of drug effects determined in ovarian cancer *DeathPro* screens in 2D and 3D culture. Dendrograms derived from hierarchical clustering of drug effects averaged over all 10 patient cell lines (AUCd=cell death, AUCpi=growth arrest, LD50=lognorm LD50, scaled to 1 for minimum and 0 for maximum dose). Subclusters are differently coloured. (**b**) 3D visualization of dendrograms shown in (a). Drug effects are averaged over all 10 patient cell lines. Drug groups derived from clustering are coloured similarly as in (a) (**c**) Differences of drug effects in patient cell lines measured with *DeathPro* assay in 2D or 3D culture after 72h (dell death - AUCd, growth arrest - AUCpi). Blue heat map colour indicates a higher drug response in 3D culture, red colour a stronger effect in 2D culture. (**d**) Differences of drug effects in patient cell lines cultured in 2D or 3D. Effects were measured with *DeathPro* assay directly after 72h drug treatment in 2D or after 72h treatment and 72h drug removal in 3D culture. Black boxes in (c) and (d) mark drugs whose effects are significantly altered in cancer organoids compared to 2D cultured cells. (**e**) Drugs whose efficiency in inducing cell death or growth arrest is significantly changed when not applied in 2D but 3D culture. Effects of drugs marked in (**c, d**). Dasatinib, AZD2014 and temsirolismus target proliferation pathways (cytostatic drugs), Azacytidine induced cell death (cytotoxic drug). HOSE=HOSEpiC, C+P = Carboplatin + Paclitaxel, *=p<0.05

A fourth drug efficacy cluster appearing in 3D, but not in 2D culture, included four drugs that induced strong growth arrest but low cell death (green cluster, Fig. 3a, b). All four drugs, ICG-001, temsirolismus, AZD5363 and AZD2014, targeted proliferation pathways and were more effective in 3D culture. To differentiate between these and other drugs, we divided our panel into ‘cytostatic drugs’ inhibiting kinases or other effectors of proliferation pathways and ‘cytotoxic drugs’ causing DNA damage, DNA methylation changes or mitotic failure.

The effects of four specific drugs and one drug combination were significantly altered in 3D compared to 2D before and after drug removal (Fig. 3c, d): Sarcoma (SRC) kinase inhibitor dasatinib induced significantly lower cell death and growth arrest in OC organoids than in monolayer patient cells (Fig. 3e). In combination with carboplatin and paclitaxel, growth arrest in organoids was still lower than in 2D. In contrast to dasatinib, the mTOR inhibitors temsirolismus and AZD2014 inhibited cell growth in organoids more strongly than in 2D culture. Azacytidine was the only cytotoxic drug that induced lower growth arrest in organoids than in cells in 2D. As azacytidine induced comparable cell death in 2D and 3D (Fig. 2c, e), its overall efficacy was similar in 2D and 3D culture (yellow, medium cytotoxic cluster Fig. 3a). In both screens we found that belinostat, BKM120 and carboplatin were the most potent drugs and that the efficacy of the cytostatic drugs dasatinib, temsirolismus and AZD2014 depended on culture type.

### Drug responses in patient organoids are more diverse and of lower therapeutic potential

Having compared drug effects generally and separately, we inspected differences and similarities of patient cell responses in 2D and 3D culture by hierarchical clustering. Interestingly, drug response profiles tended to cluster based on the patients as well as the culture format (Fig. 4a), indicating that culture type can influence patient cell responses to the same extent as intrinsic tumour heterogeneity. Most 2D patient profiles clustered together homogenously, with the exception of OC12 and OC18 which showed comparable response profiles in 2D and 3D. In total, we found 2D drug profiles in 4 subclusters while 3D drug profiles occurred in 8 subclusters, demonstrating once more that drug profiles appear more diverse in organoids. Normal HOSEpiC cells clustered separately from patient cells in 3D but showed a drug response similar to OC19 in 2D, suggesting that the culture format can conceal differences in genomic aberrations and gene expression.

**Figure 4:**
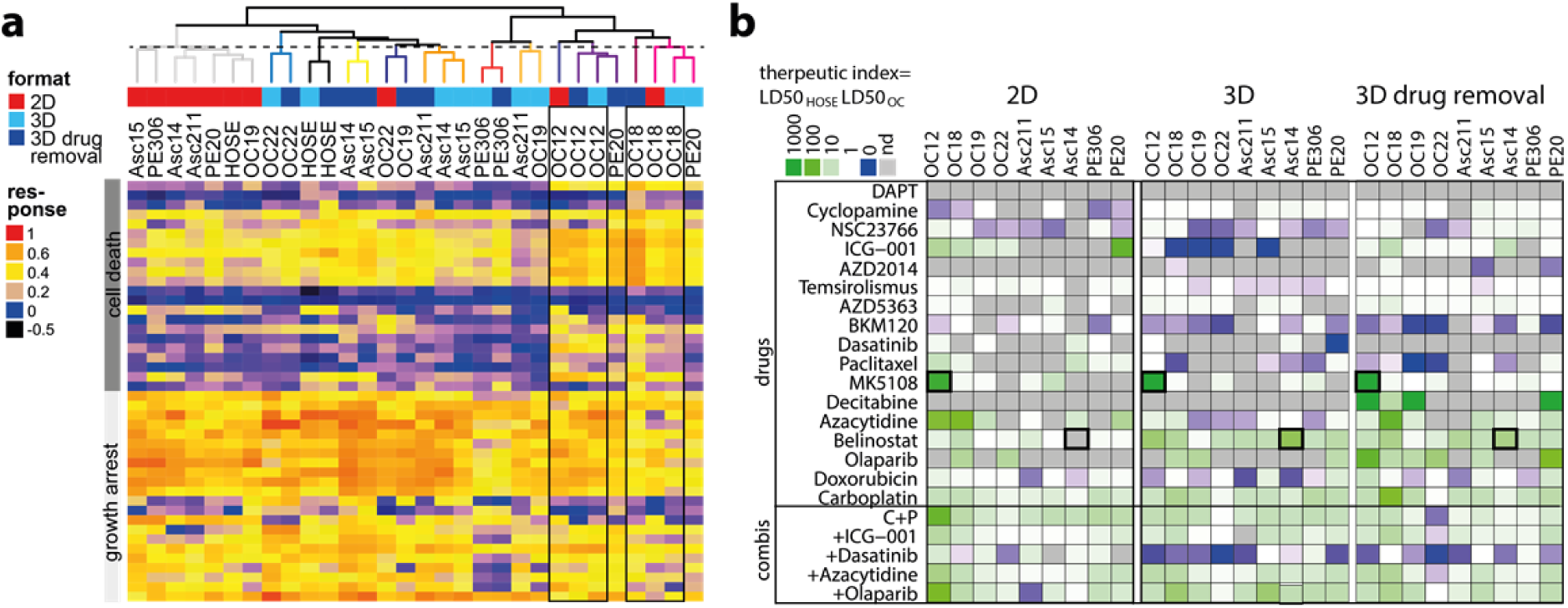
Patient organoids respond more diverse to drugs and with lower therapeutic potential than 2D cultured patient cells. **(a)** Hierarchical clustering of drug response profiles determined in ovarian cancer organoids or 2D cultured patient cells of the same origin. The dashed vertical line cuts the dendrogram arbitrarily at the height of the 2D sub cluster (grey). (**b**) Therapeutic indices determined from LD50 values derived from ovarian cancer (OC) drug screens in 2D culture or 3D culture. Green heat map colour indicates drug effectivity in cancer cells and low toxicity in normal cells, blue colour high toxicity in normal cells and low effectivity in cancer cells. nd = not determined, i.e. no fitting performed due to low response.

In our screens, we included ovarian epithelial cells (HOSEpiC) to examine cytotoxicity induced in noncancerous cells. To normalize drug efficacy in PDCLs to HOSEpiC, we calculated the therapeutic index (TI) as the ratio of LD50 values from PDCL and HOSEpiC in both cultures (Fig. 4b). TI patterns in cancer organoids were less favourable overall than in 2D culture (blue colour, Fig. 4b). The most effective candidates, carboplatin and belinostat, had positive TIs and low toxicity, while low TIs for BKM120 reflected high toxicity in normal cells. To identify patient-specific treatment options, the drug with the highest TI can be selected for each individual, e.g. MK5108 for OC12. For some patient cells, e.g. Asc14, belinostat would be suggested from organoid testing but not from 2D cell testing where cell death was too low to determine an LD50. Even if drug-induced cytotoxicity differed only minimally between 2D and 3D cultured patient cells (Fig. 2, Supplementary Fig. 5), therapeutic potentials in OC patient organoids were altered distinctively. By taking into account the heterogeneity in drug responses, our *DeathPro* assay allows the systematic deduction of patient-specific treatment options across cell cultures and patient cell lines.

### Patient cells harbour numerous copy number alterations not linked to drug-induced cell death

To predict or functionally link drug sensitivities to genetic alterations, several studies have integrated drug sensitivity data from viability assays of patient cells or cell lines with genome sequencing data^3,4,15^. Here, we performed whole genome sequencing (WGS) to associate OC genotypes with drug sensitivity data. First, we confirmed that genetic alterations in our PDCL set matched those observed in tumours: We found multiple copy number alterations (CNA) in all PDCLs (Fig. 5a), as previously reported for serous OC^16^. In a set of OC-relevant genes selected from literature^17–19^ and databases^20,21^, few deletion/insertion polymorphisms (indels) or mutations were detected, except for *TP53*, which was mutated with a similar frequency as in the COSMIC cohort (Fig. 5b). Likewise, genes frequently amplified (*MYC, PIK3CA* and *AURKA*) or lost (*RB1, PTEN*) in ovarian tumours^19^ were also commonly multiplied or lost in our set of patient cells (Fig. 5b). Since drug sensitivity frequently correlates with alterations in the corresponding drug target^15^, we associated target genes commonly affected by CNAs with cell death (AUCd) induced by the respective inhibitor. Amplifications of *AURKA* and *PI3KCA* did not alter cytotoxicity induced by AURKA inhibitor MK5108 or PI3K inhibitor BKM120, respectively (Fig. 5 c, d). Moreover, loss of *BRCA1/2*, a putative marker for impaired DNA repair capacity^22^, did not affect sensitivity towards the DNA damage related drugs carboplatin and olaparib (Fig. 5e, f).

**Figure 5:**
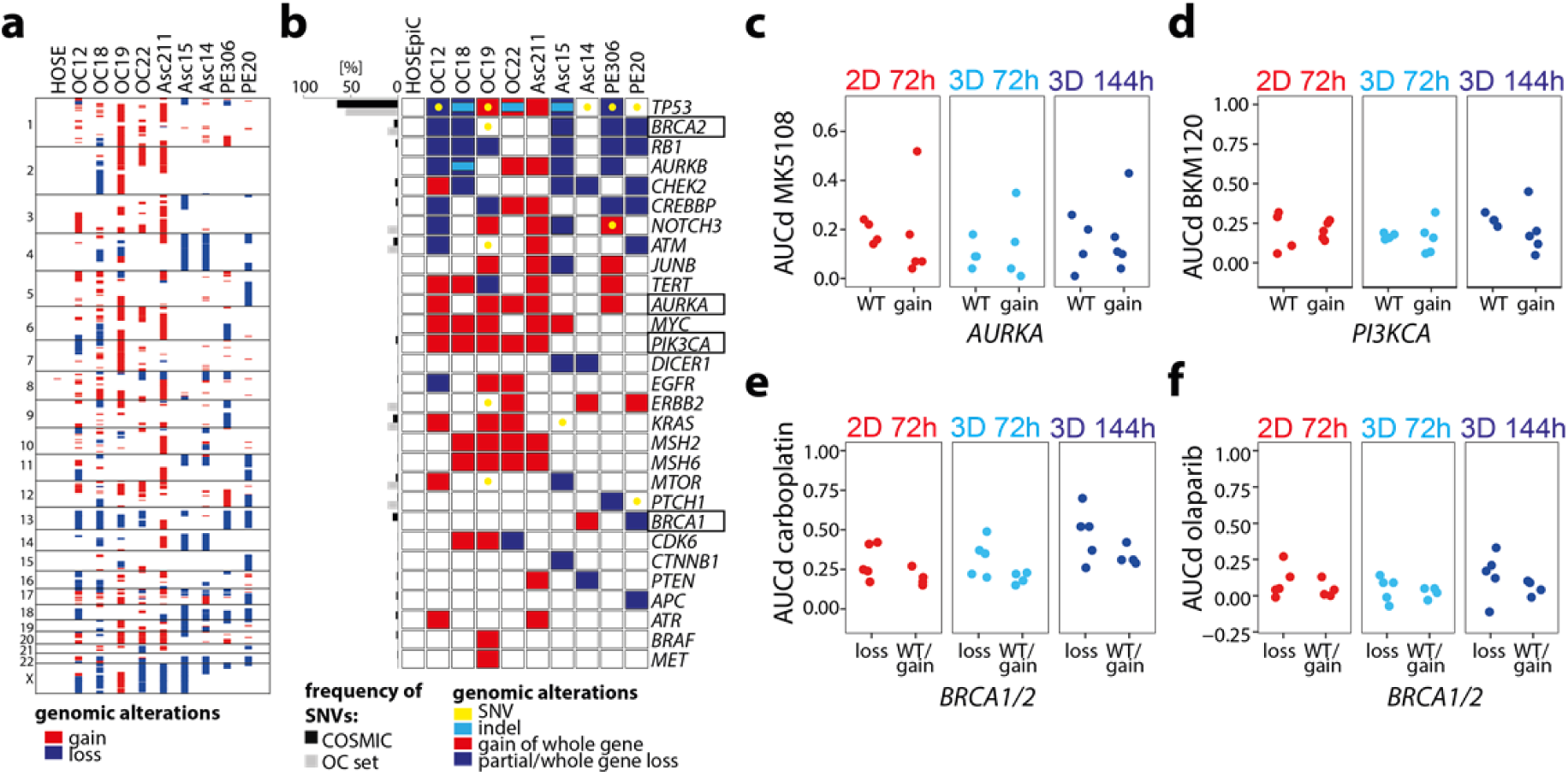
Ovarian cancer cells harbour numerous copy number alterations unlinked to drug-induced cell death. (**a**) Copy number alterations (CNAs) in primary ovarian cancer (OC) cell lines used for *DeathPro* drug screening. CNAs were determined from whole genome sequencing data. Losses are shown in blue, gains in red. (**b**) Panel of 29 OC-relevant genes depicting patient cell line-specific gains, losses including loss of heterozygosity, indels and somatic nucleotide variations (SNVs) in coding regions. Genes were selected from COSMIC and ICGC databases. (**c-f**) Association of copy number changes in drug target genes with drug sensitivities. (**c**) Cytotoxicity of Aurora Kinase A inhibitor MK5108 in patients with or without (WT) *AURKA* amplification. (**d**) Cytotoxicity of PI3K inhibitor BKM120 in patients with or without (WT) *PI3KCA* gain. (**e, f**) Cytotoxicity induced by carboplatin (**e**) or olaparib (**f**) in patients with or without (WT) *BRCA1 or BRCA2* loss.

### Homologous recombination deficiency scores correlate with drug effects in organoids

To incorporate the complex genomic aberrations in OC, we focused on the genome structure altered by DNA repair deficiencies. Loss of heterozygosity regions can be counted and added up to the homologous recombination deficiency (HRD) score (Fig. 6a) which is linked to cellular HR repair capacity^22^. HRD scores in our OC set varied between 3 and 22 (Fig. 6b). We systematically associated HRD scores and OC drug responses in different culture systems and found 20 statistically significant correlations (Rsquare >0.61, false discovery rate <0.1, Fig. 6c). Remarkably, 90% (18/20) of these potentially relevant associations were observed with 3D culture derived data. HRD scores correlated not only with cytotoxic responses to carboplatin and all its combinations (Fig. 6c, d) but also with paclitaxel, azacytidine and decitabine responses although these drugs did not directly affect DNA structure or repair (Fig. 6e, f, g). Moreover, HRD scores were linked to growth arrest induced by temsirolismus (Fig. 6d). Stratification based on high (>=10) or low (<10) HRD scores divided OC cells into responders (OC12, OC18 and PE20) and non-responders to carboplatin, olaparib or azacytidine (Fig. 6d, f, Supplementary Fig. 7a). The OC responders grew faster than non-responders in organoid but not in 2D culture (Fig. 2c, d, Supplementary Fig. 7 b, c). Thus, high HRD scores co-occurred not only with high drug-induced cytotoxicity but also with fast growth in organoids. Altogether, the strong correlation of growth and HRD scores with drug response in cancer organoids supports our view that organoids are a better model to assess patient specific drug response *in vitro*.

**Figure 6:**
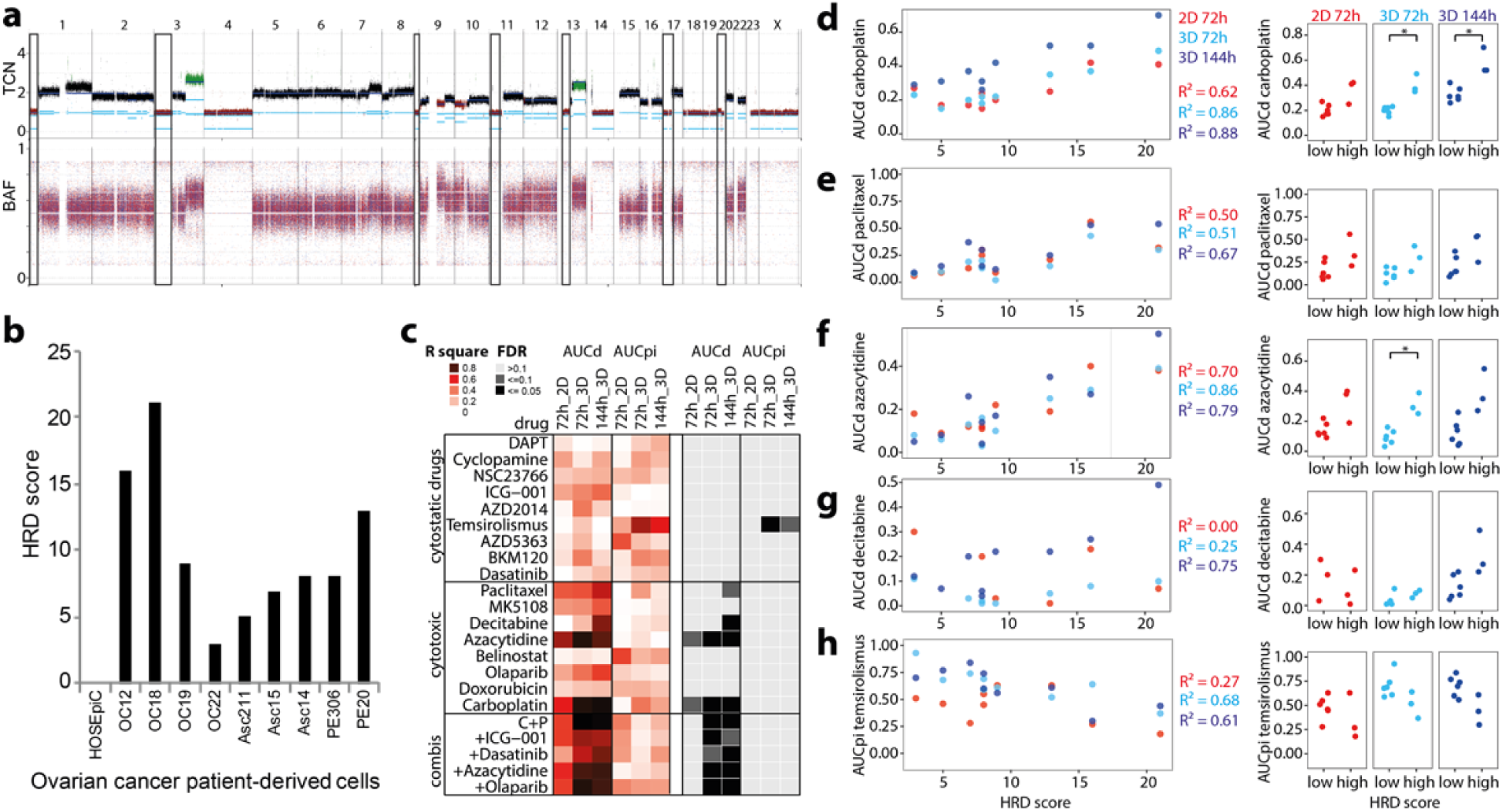
Homologous recombination deficiency scores correlate with drug-induced cell death in primary ovarian cancer cells. (**a**) Visualization of homologous recombination deficiency (HRD) score determination by counting lost chromosome regions. Total copy number (TCN) and biallelic allele frequency (BAF) plots derived from Asc15 whole genome sequencing data are shown. Black squares illustrate chromosome regions summarized as HRD score. (**b**) HRD score of patient-derived ovarian cancer cell lines used for *DeathPro* drug screening. (**c**) Heat map of correlation coefficients (R square) and estimated false discovery rates (FDR) determined from systematic association of drug responses (AUCd-cell death, AUCpi-growth arrest) with HRD scores. FDR was estimated by random sampling. (**d-h**) Drug-induced cell death (AUCd) or growth arrest (AUCpi) of all nine primary OC cell lines divided into two groups with low (<10) or high (>=10) homologous repair deficiency (HRD) score. Cytotoxicity induced by carboplatin (**d**), paclitaxel (**e**), azacytidine (**f**) and decitabine (**g**) correlates with HRD score. Growth arrest induced by temsirolismus (**h**) is reduced in HR deficient cells. *=p<0.05

## Discussion

In this study, we systematically compared drug responses between 2D and organoid cultures of patient cells and their association to genomic alterations. For this purpose we developed *DeathPro*, an automated microscopy-based workflow that simultaneously discriminates cytotoxic and cytostatic drug effects over time. Previous microscopy-based drug assays in 3D cell cultures or organoids focused on morphological changes^23,24^, metabolic parameters^25^ or required specific instrumentation to resolve cell death and growth^25,26^. Tested in a small number of cell models and not in parallel in 2D cultures, the usability and scalability of these assays is limited^24–26^. We have demonstrated the versatility and robustness of *DeathPro* in drug screens of heterogeneous OC cells in monolayer and organoid culture and in lung cancer as a second cancer entity. Unlike most image-based viability assays, which detect viable cells by cytoplasmic staining with Calcein AM^24,27^, *DeathPro* directly compares the area of nuclei of dead and live cells and generates drug efficacy measures over time independent of cellular morphology and cytoplasmic stains. Counting dead and live cells as an alternative to area measurements would require detailed, time-consuming imaging of organoids unfeasible in high-throughput drug screens. To the other end, subtle changes in nuclear size due to mitosis defects and apoptosis might be neglected. Even though we here presented drug screens based on Hoechst for staining live cells, the *DeathPro* image analysis workflow provided can be readily adapted to other nuclear stains or markers.

Resolving drug effects in OC patient cells, we found that drug-induced cell death was similar in both culture types whereas growth arrest varied. Accordingly, the efficacy of cytostatic drugs like dasatinib, temsirolismus or AZD2014 was culture type-dependent. Since most newly developed drugs are cytostatic^28^, our results highlight the importance of choosing the right model system to evaluate drug efficacy e.g. in preclinical studies. In particular, our results reveal diverse drug responses in organoids and suggest that specific drug response phenotypes are visible in organoids but not in monolayer culture. We observed less cell death in 3D compared to 2D cultures after 72h but higher death after drug removal (114h), which may lead to an underestimation of drug effects in 3D after a 72h standard treatment interval. The previously reported findings that standard cell lines in 3D culture are more chemoresistant than in 2D culture^10,29^ may therefore in part reflect altered cell death kinetics, which should be accounted for in future screens. Interestingly, the observed drug effects in OC patient cells mirrored findings from clinical trials. The combination of carboplatin and paclitaxel did not perform better than carboplatin alone, consistent with the ICON 3 trial^30^. Paclitaxel killed only 2 of 9 patient cancer organoids, similar to taxol monotherapy studies in metastatic or refractory OC that reported 20% responders^31,32^. Dasatinib, which failed at clinical phase II for recurrent OC and primary peritoneal carcinoma^33^, was effective in 2D but ineffective in 3D culture in our screen. From the drugs included in our panel, no candidate surpassed the first-line therapeutic carboplatin with regard to (i) efficacy in the whole patient set and (ii) limited toxicity in normal epithelial cells. Still, initial cytotoxicity profiles determined with *DeathPro* readily suggested patient-specific alternatives to carboplatin, such as Aurora kinase A inhibitor MK5108 for chemosensitive patient OC12 or belinostat for chemoresistant Asc14.

By associating resolved drug responses in OC patient cells with HRD scores from WGS data, we found a set of relevant correlations which would not be detected with proliferation-based assays in 2D. As expected based on a study that linked higher platinum sensitivities to HR deficiency^34^, carboplatin-induced cytotoxicity correlated with HRD scores in OC patients. Moreover, we detected correlations of DNA demethylating drug effects with HRD scores, suggesting a link between deficient DNA homologous recombination repair and DNA demethylation. While only decitabine sensitivity has been linked to *KRAS* status so far^35^, there is increasing evidence that both azacytidine and decitabine induce reactive oxygen species which cause DNA damage and finally apoptosis in cancer cells^36–38^. For all drugs whose effects correlated with HRD score, we observed a stronger correlation in OC cancer organoids than in monolayer culture. Interestingly, cell growth, HRD score and drug-induced cytotoxicity were linked in organoids but not in 2D cell culture. Similar to drug efficacy, this suggests that genotype-drug sensitivity correlations are more pronounced in 3D cultures, which is particularly important since comprehensive studies so far have focused on 2D culture data^15,39^.

Taken together, we developed and provide the *DeathPro* assay as a tool for refined drug screening and for deciphering genotype-drug sensitivity associations, and found that culture type was a key determinant of the efficacy of cytostatic drugs. In our hands, drug sensitivity was not generally decreased in organoids as previous studies suggested; instead, drug responses were more diverse and correlated better with genomic alterations in 3D compared to 2D culture. Overall, these results could provide a rationale to select the appropriate culture format for drug sensitivity assays in basic and future translational research.

## Materials and Methods

### Patient-derived cell lines and drugs

Tumour material from serous ovarian cancer patients was collected at the Departments of Gynaecology and Obstetrics, at the University Medical Centres Mannheim and Heidelberg. The study was approved by the ethical committees of the Universities of Mannheim and Heidelberg (case number 2011-380N-MA and S-008/2009) and conducted in accordance with the Helsinki Declaration; written informed consent was obtained from all patients. Primary serous ovarian carcinoma cell lines were established by transplantation of primary tumour specimen or tumour cells as previously described ^40,41^. Defined-serum-free culture medium was used as previously described ^41^ with the addition of 36 ng/ml hydrocortisone, 5 μg/ml insulin and 0.5 ng/ml beta-estradiol. HOSEpiC cells were obtained from ScienCell Research. PDCLs were checked for cross-contamination with standard OC cell lines and tested for mycoplasma contamination using the commercial Multiplex Cell Line Authentication and Mycoplasma Test Services (Multiplexion, Heidelberg, Germany). OC22 WGS data contained ∼30% reads mapping to the mouse genome due to irremovable, immortalized mouse fibroblasts potentially derived from the mouse xenograft. We used the 70% human sequences for further analysis. WGS data of all other PDCLs and HOSEpiC contained only human DNA sequences. All OC cells were cultured in Primaria flasks, subcultured using StemPro Accutase (ThermoFischer) and used at passages below 20 (PDCLs) or 6 (HOSEpiC). The cell lines LN2106 and T2427 were generated from human squamous cell carcinomas as described previously^42^. Their use for research was approved by the ethical committee of the University of Heidelberg (S-270/2001). LN2106 and T2427 cells were cultivated in DMEM/Ham’s F-12 (ThermoFischer) with 10% fetal calf serum (ThermoFischer) for not more than 20 passages. Drugs were dissolved in DMSO, water, PBS or ethanol and stored as single-use aliquots at -80 °C (table S2). Drug dilution series (1:3) were prepared using culture medium.

### *DeathPro* microscopy-based drug screen

Drug concentrations, treatment intervals and endpoints were chosen according to published studies or determined in pilot experiments. All image data were used and analysed. To assess reproducibility, the drug screens in OC cells and organoids were performed twice independently with different cell passage numbers and different drug plate layouts. Lung cancer organoids were screened once. Biological variability in all tested conditions was assessed by imaging two positions per well and no other technical replicates were included. Drug screening was performed in 96-well Angiogenesis μ-Plates from ibidi. Cells were seeded directly onto the plate for 2D culture or on growth-factor reduced, phenol red-free Matrigel (Corning, >9 mg/ml protein). Medium for drug treatment contained 1 μg/ml Hoechst (Invitrogen) and 1 μg/ml PI (Sigma) and was added one day (2D) or four days (3D) after cell seeding and substituted after 72 h by drug-free medium in 3D. Cells were exposed maximally to 1% DMSO or 1% ethanol in the highest drug concentrations and corresponding controls were included in the assay. Cells were imaged at similar positions after 0 h, 72 h and 144 h (only 3D) after start of drug treatment using a Zeiss LSM780 confocal microscope, 10x objective (EC Plan-Neofluar 10x/0.30 M27) and 405 nm and 561 nm diode lasers in simultaneous mode. Imaging was performed in an incubation chamber at 37 °C, 5% CO_2_ and 50-60% humidity using the Visual Basic for Applications macro ‘AutofocusScreen’ **^43^**.

### Image processing and drug response analysis

Image stacks were processed to maximum intensity projections (MIPs) with a custom-built macro in Fiji 2.0.0-rc-19/1.49m^44^. MIPs were uploaded and processed in our *‘DeathPro’* workflow in KNIME 3.1 (Konstanz Information Miner-*47*). Images were annotated with drugs and concentrations used and signals were extracted by thresholding. For the calculation, summarizing, clustering and plotting of values R version 3.3.2^46^ including packages drc^47^, stringr, ComplexHeatmap^48^, ggplot2^49^, reshape^50^ and RColorBrewer^51^ were used.

Hierarchical clustering with Euclidian distance and complete linkage was applied to compare PDCL-specific drug response profiles consisting of cell death (AUCd) and growth arrest (AUCpi) values measured over all drugs tested. Average linkage was used for drug response parameters averaged over all PDCLs.

### Whole Genome Sequencing and Analysis

Genomic DNA from 1x10^6^ primary cells was extracted using the DNeasy Blood & Tissue Kit (Qiagen), prepared with the TruSeq PCR free library kit (Illumina) and sequenced on a HiSeq X Ten (Illumina). Sequences were mapped to the human reference genome (build hg19, version hs37d5)^52^ using bwa-mem 0.7.8-r455^53^. The OC22 sample contained ∼30% mouse gDNA and thus had to be aligned to the hs37d5-mm10 hybrid reference sequence. Only reads mapped against hs37d5 were used for further analysis. Duplicates were marked with Picard 1.125 (https://broadinstitute.github.io/picard/). Somatic nucleotide variations and indels were called without matched control using our in-house workflow^54^, filtered^55^ and annotated with Annovar^56^. Copy number variations and loss of heterozygosity regions were determined by dedicated workflows and gains and losses were classified based on estimated ploidies. Homologous recombination deficiency scores were determined as previously described^22^. Sequence data has been deposited at the European Genome-phenome Archive (http://www.ebi.ac.uk/ega/), which is hosted by the EBI, under accession number EGAS00001002239.

### Statistical Analysis

Independent replicates refer to independent cell samples seeded, treated and imaged on different days. Differences between effects of drug combinations and single drugs were tested for statistical significance using a paired Student’s *t*-test. Differences between responses of different groups to one drug were assessed with a two-sided Welch’s *t*-test. P values <0.05 were considered statistically significant and indicated with asterisks. Pearson’s correlation coefficient (Rp) was used to describe the strength of correlation between biological replicates. Coefficient of determination (R^2^) was used to denote strength of linear relationships between area under curve values and HRD scores. False discovery rate for R^2^ was determined by random sampling.

## Acknowledgements

We thank A. Kopp-Schneider (Division of Biostatistics, DKFZ) for counseling statistical analysis, C. Dietz (Konstanz University), M. Waschow and L. Mayer (Theoretical Bioinformatics, DKFZ) for support with KNIME, B. Burwinkel (Molecular Epidemiology, DKFZ) for provision of several ovarian cancer samples, D. Huebschmann, G. Warsow (Theoretical Bioinformatics, DKFZ) for assistance in WGS data analysis, C. Klein and V. Vogel for immunohistochemistry and T. Krieger (Theoretical Bioinformatics, DKFZ) for critically revising the manuscript. We thank the High Throughput Sequencing unit and the Microarray unit (Genomics & Proteomics Core Facility, DKFZ) for providing whole genome sequencing and microarray services. Tissue samples were provided by the tissue bank of the National Center for Tumour Diseases (NCT, Heidelberg, Germany) in accordance with the regulations of the tissue bank and the approval of the ethics committee of Heidelberg University. The authors would like to thank the Exome Aggregation Consortium and the groups that provided exome variant data for comparison. A full list of contributing groups can be found at http://exac.broadinstitute.org/about.

## Supplementary Materials

**Supplementary Figure 1:**
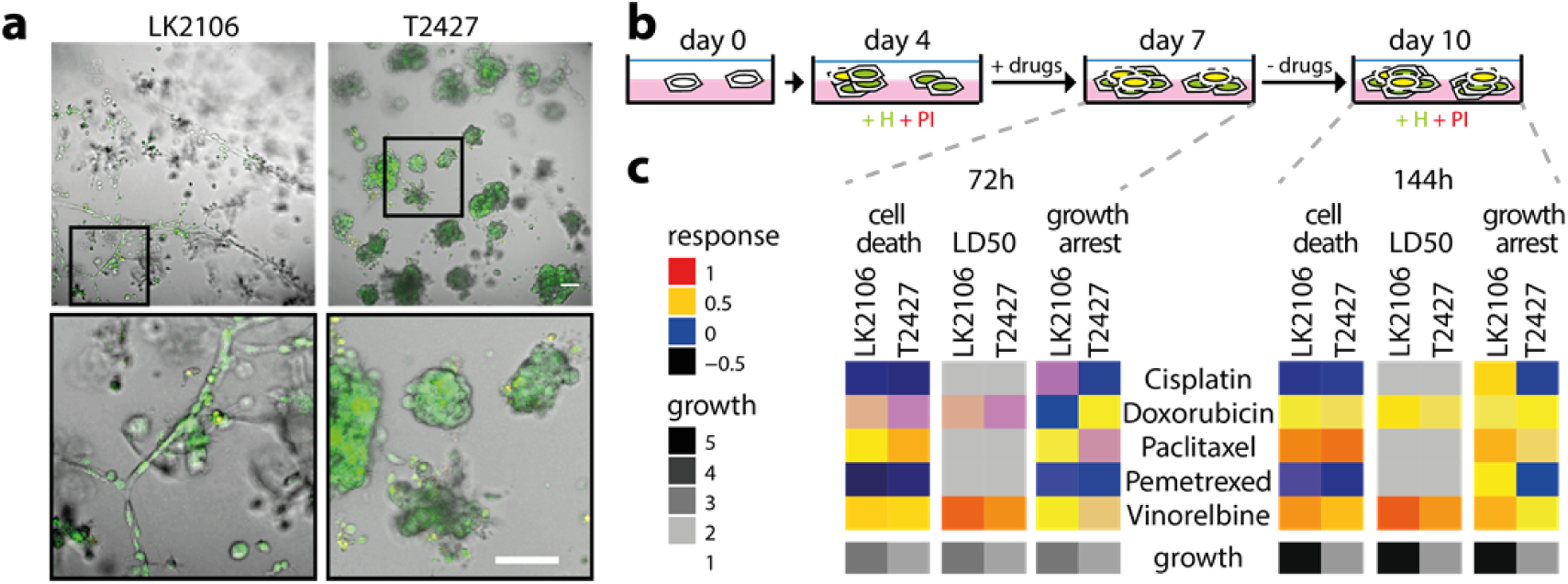
DeathPro Screen in primary cells derived from lung cancer patients. (**a**)Lung cancer cells lines derived from lymph node (LN2106) or lung tumour (T2427) were cultured on Matrigel for 7 days and stained with Hoechst (H, green) and propidium iodide (PI, red). Scale bar is 100 μm. (**b**) Overview of drug screen schedule. (**c**) Drug responses and cell growth measured after 72 h or 144 h for drugs as indicated. For better visualization, logarithmic LD50 values were normalized so that 1 and 0 correspond to minimum and maximum dose, respectively.

**Supplementary Figure 2:**
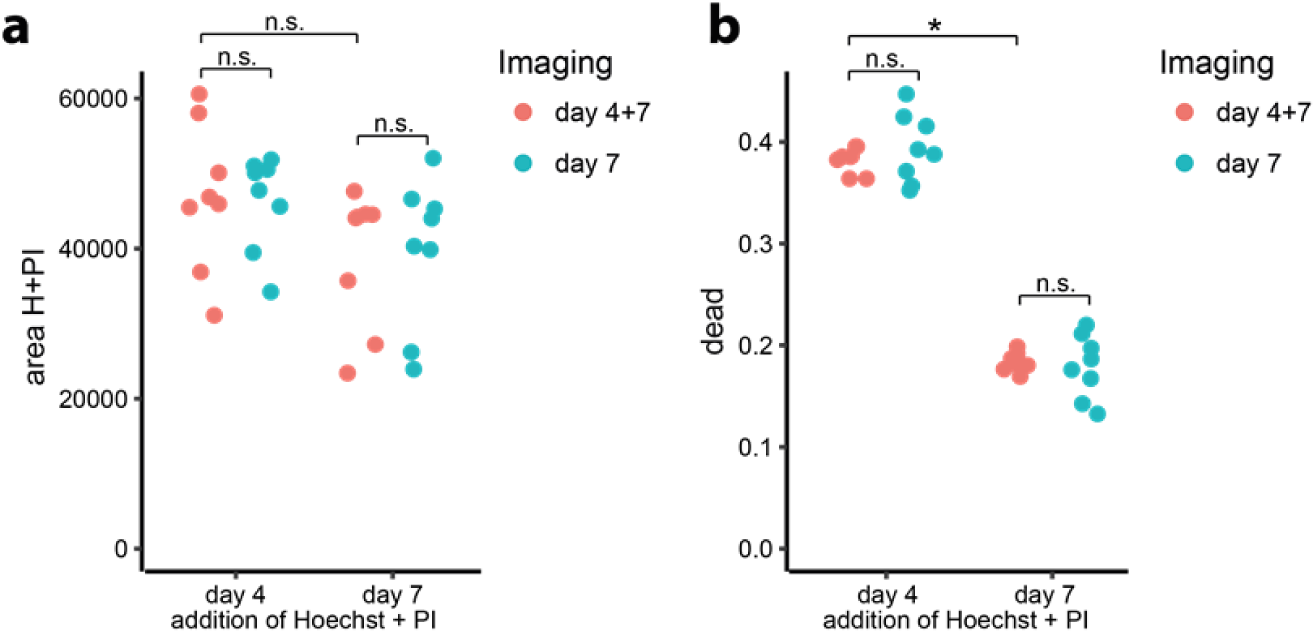
Effect of imaging conditions and dyes on cell growth and cell death. OC12 cells stained with Hoechst and PI at day 4 or day 7 after seeding and imaged once (day7) or twice (day 4+7). Values derive from one experiment, 8 images were acquired in two different wells (technical replicates). n.s.= not significant, *p-value<0.05

**Supplementary Figure 3:**
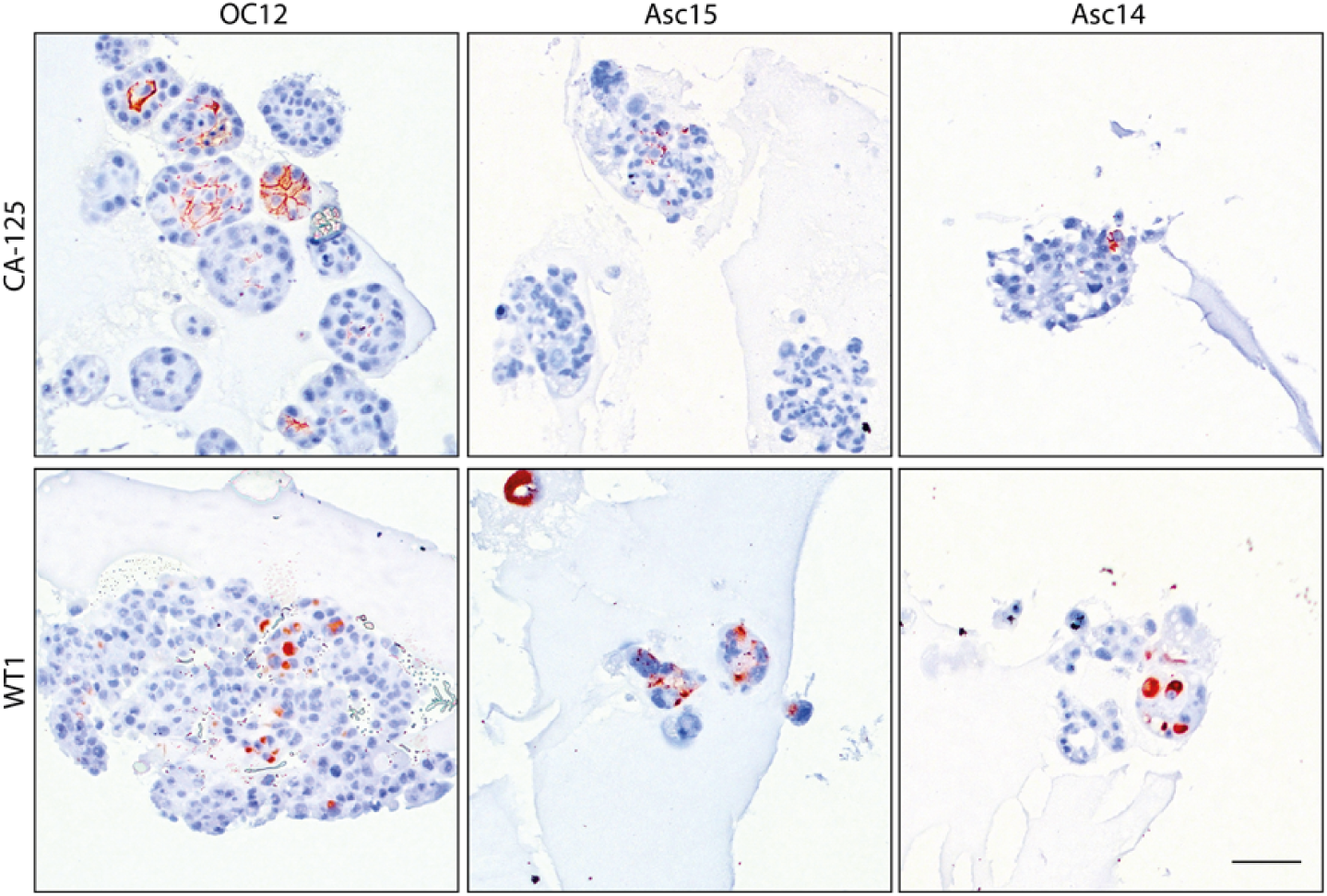
Cancer organoids from patient derived OC cells retain tumour markers CA-125 and WT1. Immunohistochemistry for CA-125 and WT1 was performed with OC12, Asc15 and Asc14 cells grown for 8 days on Matrigel. Scale bar is 200 μm.

**Supplementary Figure 4:**
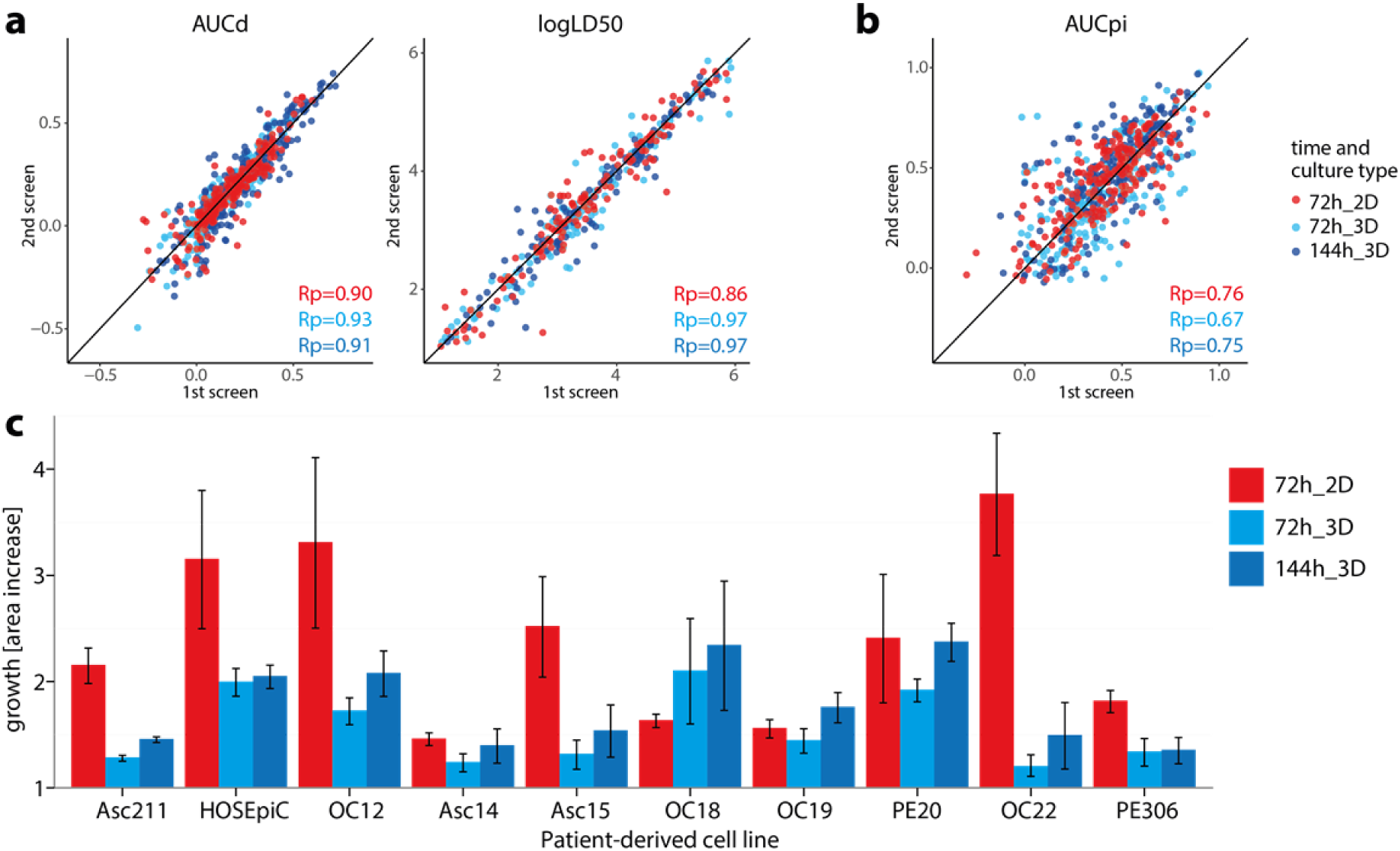
Reproducibility of DeathPro drug screens. (**a, b**) Scatterplots of AUCd, logLD50 and AUCpi values of biological replicates determined in 2D and 3D culture screens of patient-derived ovarian cancer cell lines and HOSEpiC (220 measurements per time point: 22 drug dilution series on 10 cell lines) with corresponding Pearson correlation coefficients (Rp). The black line (x=y) is depicted as reference for perfect correlation. (**c**) Growth of patient-derived ovarian cancer cell lines and HOSEpiC on Matrigel within 72 h or 144 h from day 4 on (3D) or as cell monolayers (2D) from day 1 to day 4 after seeding.

**Supplementary Figure 5:**
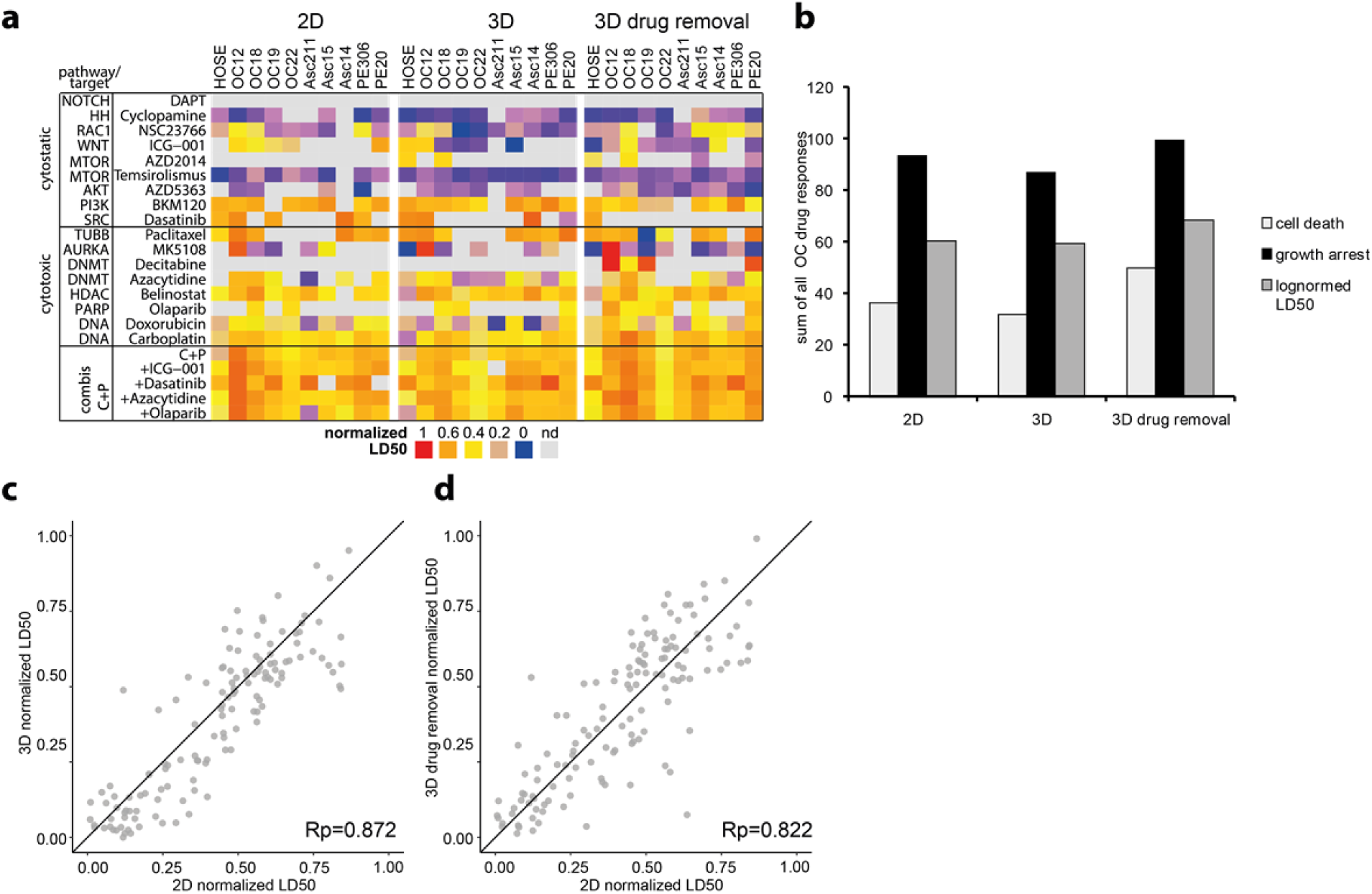
Drug sensitivity described by LD50 is similar in cells cultured in 2D or as organoids. (**a**) LD50 values determined in after 72 h drug exposure, or 72 h drug removal in OC patient derived cells cultured in 2D or as organoids. For better visualization, LD50s have been rescaled to values between 0 and 1, representing maximum and minimum dose. (**b**) Sum of all responses across all drugs and combinations and OC cells tested in 2D and 3D culture. (**c**) Comparison of LD50s in 2D vs 3D. (**d**) Comparison of LD50s in 2D vs 3D after drug removal. All values shown are means of two independent biological replicates. Rp = Pearsons correlation coefficient. C+P = Carboplatin + Paclitaxel.

**Supplementary Figure 6:**
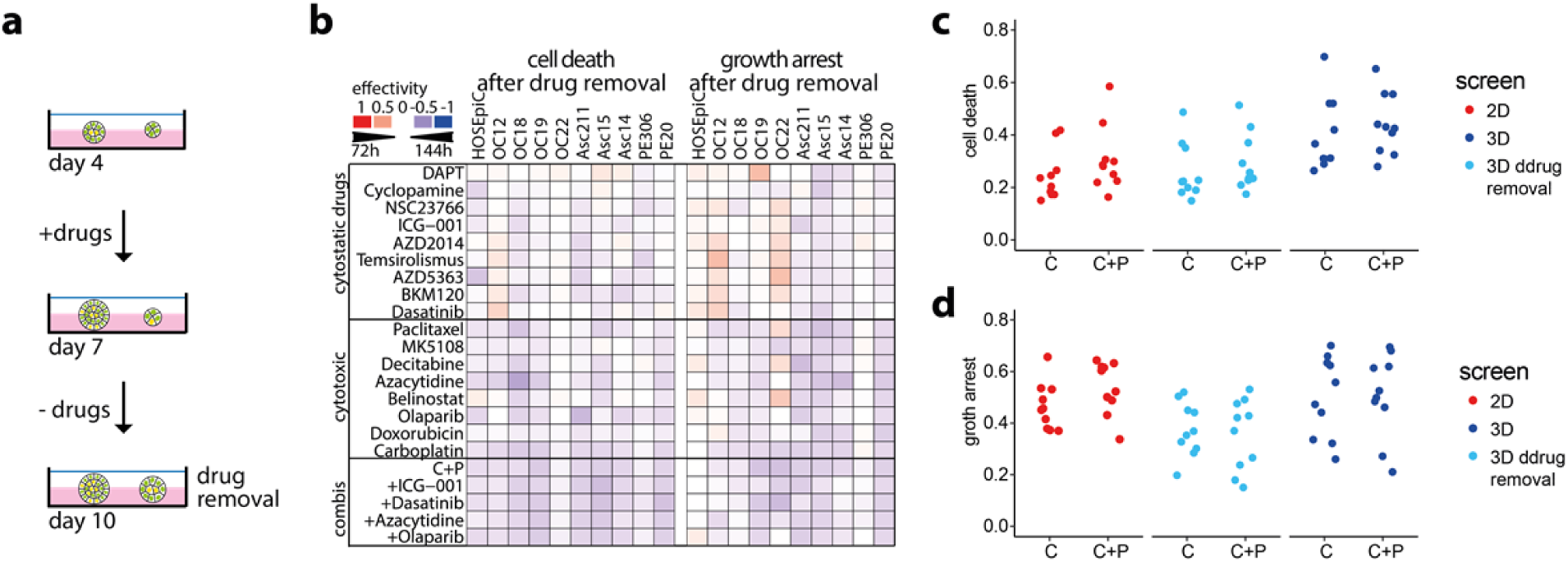
Drug-induced growth arrest and cell death increase in ovarian cancer organoids with time. (**a**) Schematic overview of drug testing in organoid culture with the DeathPro assay. (**b**) Differences of drug effects in *DeathPro* screens in OC cells over time. Cell death and growth arrest were determined after 72 h or 144 h and subtracted from each other. Blue heat map color indicates stronger effect after 144 h, red color stronger effect after 72 h. (**c**) Cell death (AUCd) induced by Carboplatin (C) or Carboplatin and Paclitaxel (C+P) in OC cells and HOSEpiC grown in 2D or 3D culture. (**d**) Growth arrest (AUCpi) induced by Carboplatin (C) or Carboplatin and Paclitaxel (C+P) in OC cells and HOSEpiC.

**Supplementary Figure 7:**
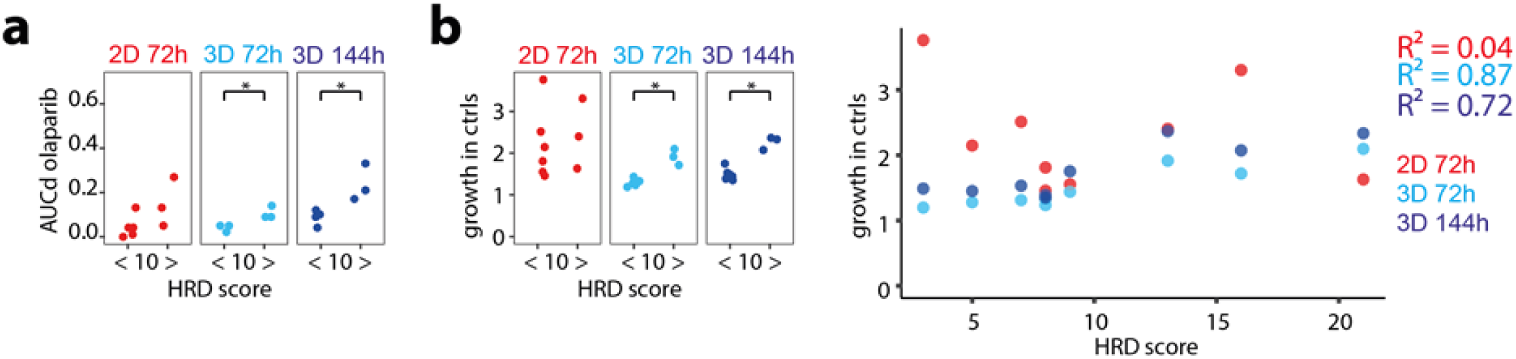
Homologous recombination deficiency scores and cell growth. (**a**) Drug-induced cell death (AUCd) of all nine primary OC cell lines divided into two groups with low (<10) or high (>=10) homologous repair deficiency (HRD) score. Cytotoxicity induced by olaparib (**a**) is higher in HR deficient cells. (**b**) Growth in untreated patient-derived OC cells correlates with HRD score in 3D culture but not in 2D culture. *=p<0.05

**Supplementary Table 1:** Classification of OC patient samples used for serous OC cell lines.

| PDCL | origin | FIGO | grade | TNM | Pathological disease | treatment |
| --- | --- | --- | --- | --- | --- | --- |
| OC12 | tumour | IIIc | G3 | TN1M1 | serous adenocarcinoma | not treated |
| OC18 | tumour | IIIc | G3 | TN1M1 | serous adenocarcinoma | not treated |
| OC19 | tumour | IIIc | G3 | TN1M1 | serous adenocarcinoma | not treated |
| OC22 | tumour | IIIc | nd | nd | serous adenocarcinoma | not treated |
| Asc211 | ascites | IIIC | G3 | pT3c<br>pN1 | serous adenocarcinoma | 1st line, 2nd line, 3rd line |
| Asc14 | ascites | IV | G3 | TN1M1 | serous adenocarcinoma | 1st, 2nd, 3rd line |
| Asc15 | ascites | IIIc | G3 | TN1M1 | serous adenocarcinoma | 1st, 2nd line |
| PE306 | pleural effusion | IV | G2 | T3 | serous adenocarcinoma | 1st line |
| PE20 | pleural effusion | IIIc | G3 | TN1M1 | serous adenocarcinoma | 1st, 2nd line: Morab study |

**Supplementary Table 2:** Inhibitors used for the *DeathPro* screens

| Inhibitors | Status in OC therapy | producer | Lot # | stock conc. [mM] | starting conc. [µM] | solvent |
| --- | --- | --- | --- | --- | --- | --- |
| paclitaxel | 1<sup>st</sup> line | Selleckchem | 9 | 10 | 1 | DMSO |
| carboplatin | 1<sup>st</sup> line | Cayman Chemical | 0453486-12 | 20 | 2,000 | water |
| doxorubicin | 2<sup>nd</sup> line | StressMarq Biosciences | 150120 | 1 | 10 | PBS |
| olaparib | 4<sup>th</sup> line | Cayman Chemical | 0461572-5 | 10 | 100 | DMSO |
| BKM120 | phase I | Selleckchem | S224704 | 50 | 100 | DMSO |
| MK-5108 | phase I | Selleckchem | S277001 | 10 | 100 | DMSO |
| belinostat | phase II | BioVision | 9C242480 | 50 | 100 | DMSO |
| AZD2014 | phase II | Selleckchem | 1 | 50 | 5 | DMSO |
| AZD5363 | phase II | Selleckchem | S801901 | 50 | 500 | DMSO |
| temsirolimus | phase II | Sigma | 110M4716V | 10 | 40 | DMSO |
| azacytidine | phase II | Sigma | MKBR7212 | 50 | 100 | DMSO |
| decitabine | phase II | Sigma | MKBR6437V | 10 | 500 | water |
| dasatinib | phase II | LC Laboratories | BDS109 | 100 | 10 | DMSO |
| cyclopamine | preclinical | Adipogen | A00152 | 4.5 | 45 | ethanol |
| DAPT | preclinical | Sigma | 014M4609V | 25 | 100 | DMSO |
| NSC23766 | preclinical | Sigma | 044M4761V | 10 | 1,000 | water |
| ICG-001 | preclinical | Selleckchem | 2 | 50 | 100 | DMSO |

